# Short repeat RNA reduces cytotoxicity by preventing the aggregation of TDP-43 and its 25 kDa carboxy-terminal fragment

**DOI:** 10.1101/2022.07.03.498631

**Authors:** Ai Fujimoto, Masataka Kinjo, Akira Kitamura

**Affiliations:** Laboratory of Molecular Cell Dynamics, Faculty of Advanced Life Science, Hokkaido University, N21W11, Kita-ku, Sapporo 001-0021, Japan; Graduate School of Life Science, Hokkaido University, N10W8, Kita-ku, Sapporo 060-0810, Japan; PRIME, Japan Agency for Medical Research and Development, Chiyoda-ku, Tokyo 100-0004, Japan

## Abstract

TAR DNA/RNA-binding protein 43 kDa (TDP-43) proteinopathy is a hallmark of neurodegenerative disorders such as amyotrophic lateral sclerosis, in which cytoplasmic aggregates containing TDP-43 and its C-terminal fragments, such as TDP25, are observed in degenerative neuronal cells. However, few reports have focused on small molecules that can reduce their aggregation and cytotoxicity. Here, we show that short RNA repeats of GGGGCC and AAAAUU are aggregation-suppressors of TDP-43 and TDP25. TDP25 interacts with these RNAs, as well as TDP-43, despite the lack of major RNA-recognition motifs, using fluorescence cross-correlation spectroscopy. Expression of these RNAs significantly decreases the cells harboring cytoplasmic aggregates of TDP-43 and TDP25 and ameliorates rounded and shrinking cells by mislocalized TDP-43; furthermore, the cellular transcriptome is not altered. Consequently, these RNAs can maintain proteostasis by preventing aggregation of TDP-43 and TDP25.

## Introduction

TAR DNA/RNA-binding protein 43 kDa (TDP-43) is the key focus of research to understand neurodegenerative diseases such as amyotrophic lateral sclerosis (ALS) and frontotemporal dementia (FTD) [1,2]. A hallmark of these disorders is cytoplasmic aggregates of TDP-43 and its carboxy-terminal fragments (CTFs). TDP-43 carries two RNA/DNA-recognition motifs (RRM1 & 2), each of which contain two highly conserved short sequence motifs required for nucleic acid binding [3], and a C-terminal glycine-rich and intrinsically disordered region (GRR). Many of the amino acid substitutive mutations in TDP-43 associated with ALS and FTD are conserved in the GRR [4]. TDP-43 is mainly localized in the nucleus but is also a nuclear-cytoplasmic shuttling protein, which maintains cellular homeostasis regulated by the quality of RNA such as mRNA and small RNA [5]. A 25 kDa TDP-43 CTF (TDP25; amino acids 220–414), which lacks RRM1 and almost half of RRM2 [6] and is produced by caspase or calpain proteases [7–9], is highly prone to aggregation and leads to cell death. TDP25 aggregates sequestrate full-length TDP-43; therefore, loss-of-function of TDP-43 may be associated with cytotoxicity in addition to gain-of-function of these aggregates [10]. In contrast, aggregates of TDP-43 and TDP25 that can be viewed by a microscope and contain amyloid-like fibrils [11,12] in the cytoplasm may not necessarily be essential for neuronal degeneration in ALS [13,14]. Mislocalization of TDP-43 contributes to the cytotoxicity in ALS through the regulation of various cellular functions such as gene expression, mRNA maturation, repression of alternate splicing, cryptic exon splicing, and alternate polyadenylation [15,16]. The long-term mislocalization of TDP-43 in the cytoplasm produces its cytoplasmic aggregates, leading to the disruption of physiological function (e.g., sequestration of cellular proteins such as other ALS-associated and autophagy regulating protein, SQSTM1); thus, the homeostatic level (proteostasis) [17,18] regulated by proteins such as SQSTM1 could be declined [19]. Thus, motor neuron degeneration may be involved in the mislocalization of TDP-43 with the formation its parallel aggregate [14].

RNA-binding proteins (RNP), such as TDP-43, and RNA interact and induce condensation of RNPs through phase separation [20]. In contrast, RNA degradation induces aggregation of TDP-43 and TDP25 [8,21]. Binding of TDP-43 to RNA antagonizes aggregation and phase transitions [22,23]. Furthermore, the idea that RNAs such as ribosomal RNA, transfer RNA, and so on, might have chaperone activity in vitro has been proposed [24–26]. Therefore, RNA regulates the aggregation and condensation of RNPs with their phase transition and plays a role in the regulation of the intermolecular assembly using the physicochemical properties of RNA. However, which RNA regulates such aggregation of ALS/FTD-causative TDP-43 and then plays a role in cytoprotection remains unclear. Here, we show that GGGGCC and AAAAUU hexanucleotide repeat RNAs (r(G4C2) and r(A4U2), respectively) can reduce proteotoxicity by preventing the aggregation of TDP-43 and TDP25 in a cellular model. The C-terminal domain of TDP-43 was found to recognize the structure of G-quadruplex (Gq) in RNA [27], and Gq DNA protect the aggregation of denatured proteins [28]. Therefore, we first focused on whether TDP25 binds to the GGGGCC hexanucleotide repeat RNA, r(G4C2), which can be folded as a Gq and is included in a major ALS-causative transcript, *C9orf72*. Then, we demonstrate whether non-ALS-associated and short r(G4C2) plays a cytoprotective role in TDP25 aggregation and mislocalized TDP-43 with its aggregation. Furthermore, we showed that the AAAAUU repeat RNA, r(A4U2), which is generally used as a control sequence for r(G4C2) [29,30], also binds to TDP-43 and TDP25, and suppresses their aggregation, in addition to reducing the cytotoxicity caused by their mislocalization and aggregation. The following results suggest that these repeat RNAs may have chaperone-like activities that protect protein aggregation and maintain intracellular proteostasis.

## Results

### GGGGCC and AAAAUU hexanucleotide repeats bind to TDP-43 and TDP25 in cell lysates

To analyze the interaction between TDP25 and RNA, we employed a single molecule detection method, fluorescence cross-correlation spectroscopy (FCCS), which can quantitatively determine the strength of interaction between diffuse fluorescent species in solution from the fluorescence fluctuation that occurs by passing fluorescent molecules through the confocal detection volume (Figure 1a) [31,32]. After mixing RNA labeled with a chemical near-infrared fluorescent dye, Alexa Fluor 647 (AF), in lysates of murine neuroblastoma Neuro2a cells expressing green fluorescent protein (GFP)-tagged TDP25 (T25), GFP-tagged TDP-43 (T43), or GFP monomers as the tag, FCCS was performed. As we have previously shown [8], the relative cross-correlation amplitude (RCA) of the AF-labeled 24-mer of uracil guanine repeat RNA (UG) with T43 was high, whereas those with GFP and T25 were not (Figure 1b), indicating that the interaction between T43 and UG as a positive control could be determined using FCCS. The RCA when AF-labeled four repeats of r(G4C2) (G4) in a coexisting state of the hairpin (HP) and Gq structures (Gq + HP) [30] was mixed with both T43 and T25 was significantly positive; this suggests that G4 can interact with not only T43 but also T25. The RCA of T43 and T25 with only HP-formed G4 was also positive (Figure 1b), suggesting that the structure of Gq may not be required for the interaction with T43 and T25. Next, to investigate whether T25 also interacts with other RNAs that are not different from G4, we tested AF-labeled four repeats of r(A4U2) (A4), which is often used as a control sequence for r(G4C2) [29,30]. The RCA of T43 and T25 with A4 was positive but lower than that with G4 in a mixed state of the hairpin (HP) and Gq structures (Figure 1b), suggesting that T43 and T25 may be able to interact with G4 and A4, regardless of their steric structures.

**Figure 1.**
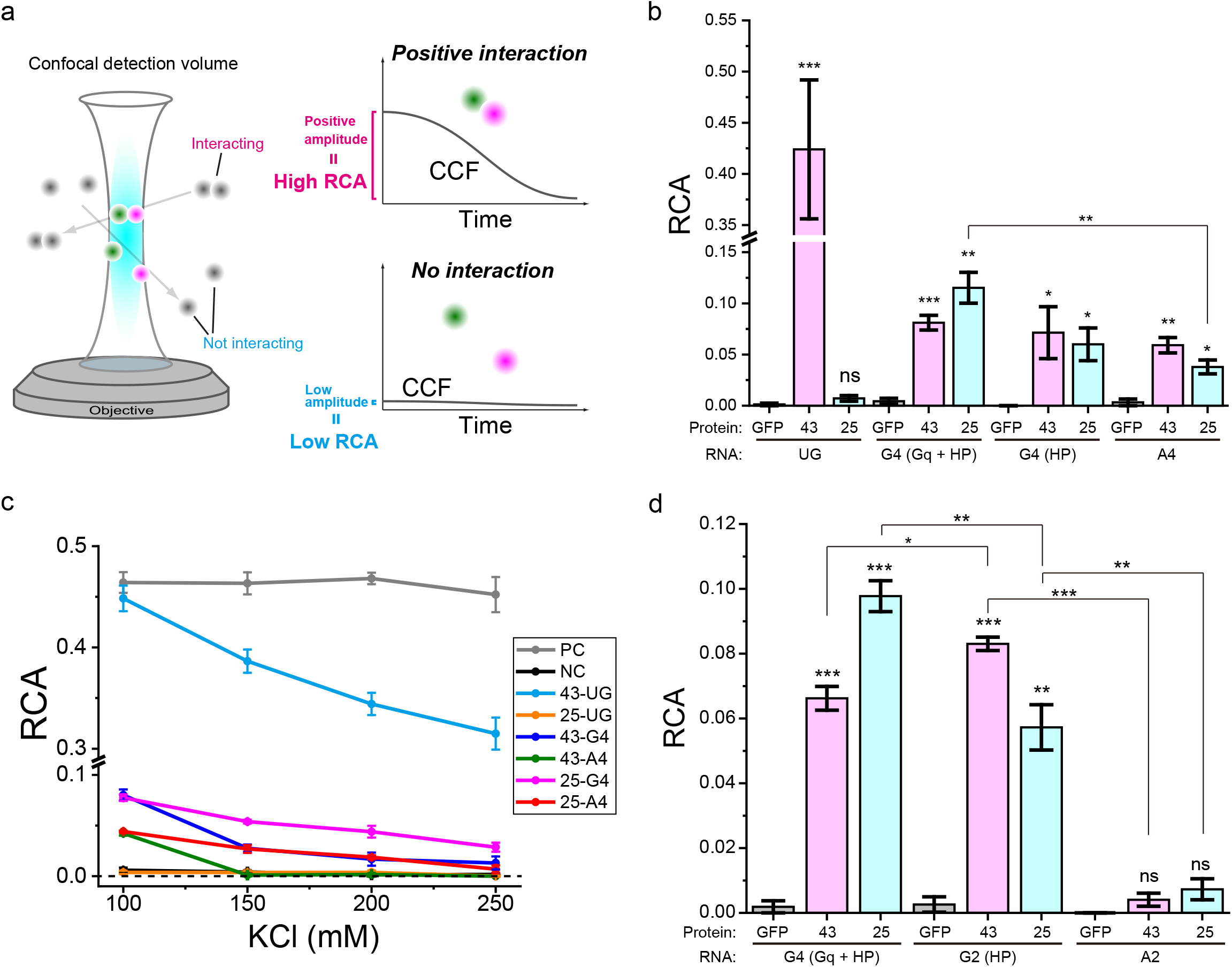
Interaction of the G4C2/A4U2-repeat RNA with TDP-43 and TDP25 in cell lysates using fluorescence cross-correlation spectroscopy. a) Illustration of the detection of molecular interactions from the relative cross-correlation amplitude (RCA) using fluorescence cross-correlation spectroscopy. b) RCA values when AF-labeled RNAs (UG, G4, and A4) that were folded in 100 mM KCl (Gq + HP) or 100 mM LiCl (HP) were mixed in lysates of Neuro2a cells expressing GFP monomers (GFP), GFP-TDP-43 (43), and GFP-TDP25 (25) with 100 mM KCl (n = 3; dots and bars indicate independent trials and mean ± SE). c) RCA values between AF-labeled RNAs (UG, G4, and A4) and GFP-tagged TDP-43/TDP25 in cell lysates at 100, 150, 200, and 250 mM KCl concentrations. G4 (Gq + HP) was folded in the presence of 100 mM KCl, then mixed with the cell lysate. The dashed line indicates zero. d) RCA values when AF-labeled RNAs (G4, G2, and A2) were used. G4 (Gq + HP) was folded in the presence of 100 mM KCl, then mixed with the cell lysate including 100 mM KCl. G2 (HP) was folded in presence of 100 mM LiCl, then mixed with the cell lysate including 100 mM KCl (n = 3; mean ± SE). (b & d) Significance: \**p* < 0.05, \*\**p* < 0.01, \*\*\**p* < 0.001, and ns (*p* ≥ 0.05).

Next, to demonstrate biochemical properties in the interaction between T43/T25 and G4/A4, the interaction strength in presence of high concentrations of KCl (150, 200, and 250 mM) was analyzed using FCCS. The RCA of neither dual color-fluorescent DNA [33] as a positive control nor the mixture of the dual fluorescent dyes as a negative control changed at the high concentrations of KCl compared to that at 100 mM (gray and black lines in Figure 1c), indicating that the FCCS detection system did not change at high KCl concentration. The RCA between T43 and UG little by little declined but not completely as the increase in KCl concentration (cyan line in Figure 1c), suggesting the strong electrostatic interaction between them. The RCA value between T25 and UG was not completely detected at all KCl concentrations (orange line in Figure 1c). The RCAs between T43 and G4/A4 prominently declined with the increase in KCl concentration (blue and green lines in Figure 1c); however, those between T25 and G4/A4 did not precipitously decline with the increase in KCl concentration (pink and red lines in Figure 1c). The RCA between T25 and G4/A4 was still positive and higher than that between T43 and G4/A4 at 150 and 200 mM KCl concentrations (Figure 1c). Therefore, the interaction between TDP25 and G4/A4 may not only be electrostatic but also hydrophobic; however, the electrostatic interaction may relatively contribute to that between TDP-43 and G4/A4.

Next, to determine whether the shorter repeats are required for this interaction, two repeats of r(G4C2) (G2) that were folded in presence of 100 mM LiCl and not KCl, [34] and two repeats of A4U2 RNA (A2) were used. Although the positive RCAs were observed between T43/T25 and AF-labeled G2, they were lower than that with G4 (Figure 1d). However, the RCA between T43/T25 and AF-labeled A2 was not observed (Figure 1d and Supplemental Figure S1). These results suggest that the chain length dependency of the interaction strength between TDP-43/TDP25 and r(G4C2) and r(A4U2), and the interaction of r(G4C2) with TDP-43/TDP25 could be weaker than that of r(A4U2). Furthermore, the Gq structure may not be necessarily required for this interaction. Although 4–23 repeats of G4C2 in the *C9orf72* transcript are associated with the formation of a cellular condensate—granule of RNP—more than 23 repeats lead to ALS/FTD-associated phase transitions of RNP, such as TDP-43, because the multiple and tandem Gq may work as a scaffold of RNP assembly [35,36]. Because the stable and abundant Gq structure can likely lead to such phase transitions, the binding ability of TDP-43 and TDP25 not only to Gq but also to the HP structure may be an important physicochemical aspect for the regulation of their aggregation/condensation.

### Direct interaction between GGGGCC/AAAAUU hexanucleotide repeats and TDP-43/TDP25

Because the above analysis was performed in a cell lysate, which is an advantage of FCCS because the biomolecular interaction can be measured in a solution mixture containing fluorescent molecules, it is not yet clear whether T25 interacts with RNA directly or cellular RNA-binding proteins in complexes of T25 interact with RNA. To confirm the direct interaction with these RNAs, we prepared purified GFP, T43, and T25 through tandem affinity purification using polyhistidine and strep tags. Sodium dodecyl sulfate polyacrylamide gel electrophoresis (SDS-PAGE) followed by silver staining showed that highly purified GFP was obtained; however, several cellular proteins were included in the purified T43 and T25 samples (Figure 2a). Since T43 is an RNP and binds to RNA with Gq-foldable sequences [27,37], we identified the co-purified proteins with T25 using peptide mass fingerprinting with mass spectroscopy (PMF-MS). Four of the five identified major proteins co-purified with T25 were not RNP (pyruvate carboxylase, tubulin, and actin by PMF-MS, and a fragment containing GFP by western blotting), while HSP70, which has been reported to have RNA-binding properties [38], was identified (Figure 2a). To confirm whether HSP70 that was co-purified with T25 bound to G4/A4, AF-labeled G4/A4 was mixed in the lysate of cells expressing HSP70 tagged with GFP, and following FCCS analysis under the same conditions as T43/T25 did not show their significant interaction (Supplemental Figure S3). Since HSP70 is an abundant molecular chaperone and is frequently co-precipitated, the interaction with HSP70 may contribute to maintaining the solubility of T25. In other words, it may be difficult to recover T25 without HSP70 in soluble fraction. Because the endogenous TDP-43 band was not observed in the purified T25 sample (Figure 2a), and the interaction between T25 and TDP-43 was not detected using FCCS in cell lysate [8], it is unlikely that the T25 sample contains endogenous TDP-43. Consequently, other RNPs may not be included in the purified T25. The interaction of purified GFP, T43, and T25 with G4 or A4 was then analyzed using FCCS. No positive RCA of the GFP, T43, and T25, when mixed with just AF dye, was observed; however, positive RCA of T43 and T25, when mixed with G4 and A4, was observed, and it was higher with G4 than with A4 (Figure 2b). Consequently, T43 and T25 interact directly with G4 and A4, and not through other RNPs.

**Figure 2.**
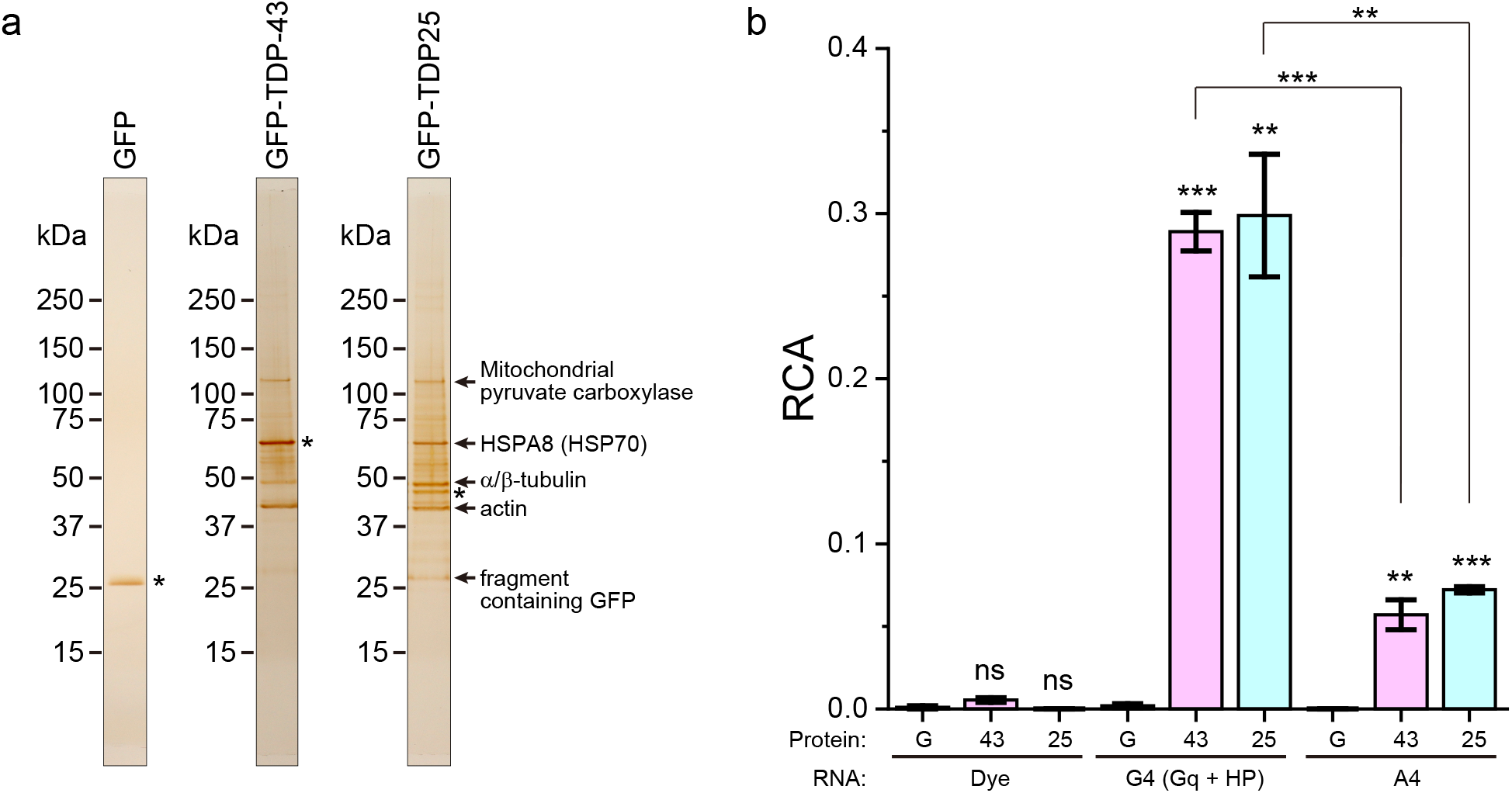
Direct interaction of the G4C2/A4U2-repeat RNA with purified TDP-43 and TDP25. a) SDS-PAGE of purified GFP, GFP-TDP-43, and GFP-TDP25, followed by silver staining. Asterisks show purified target proteins. b) RCA values when AF-labeled RNAs (G4, and A4) folded in 100 mM KCl were mixed with purified GFP monomers (GFP), GFP-TDP-43 (43), and GFP-TDP25 (25) with 100 mM KCl (n = 3; mean ± SE). Significance: \*\**p* < 0.01, \*\*\**p* < 0.001, and ns (*p* ≥ 0.05). The typical normalized cross-correlation functions are represented in Supplemental Figure S2.

### Truncated RRM is responsible for the interaction between GGGGCC/AAAAUU hexanucleotide repeats and C-terminal fragments of TDP-43

How does T25 recognize RNA even though it lacks many of the RRMs? To identify the contribution of truncated RRM in TDP-43 CTFs to the interaction with G4/A4, the interactivity of two additional truncated variants of TDP25 (C233 and C274 [39]; Figure 3a) with G4/A4 was analyzed using FCCS in cell lysates. Although C233 entirely lacks the highly conserved short motifs required for nucleic acid binding [3], C233 interacted with G4 in a mixture of Gq and HP, HP-fold G4, and A4; in contrast, C274, just the GRR, did not (Figure 3b), suggesting that the truncated RRM region may be responsible for the interaction with RNA not through the highly conserved nucleic acid binding sequences. Furthermore, the RCA between C233 and G4 with the HP fold was decreased compared to that in a mixture of Gq and HP of G4 (Figure 3b). Although TDP25 forms hetero-oligomers with cellular proteins, C233 preferentially forms homo-oligomers [39]. Because the differences in the components in the oligomers would affect their structure, such structural differences likely contribute to the recognition of Gq-RNA. Consequently, these analyses identified that oligomeric states of TDP25 would recognize G4/A4-RNA.

**Figure 3.**
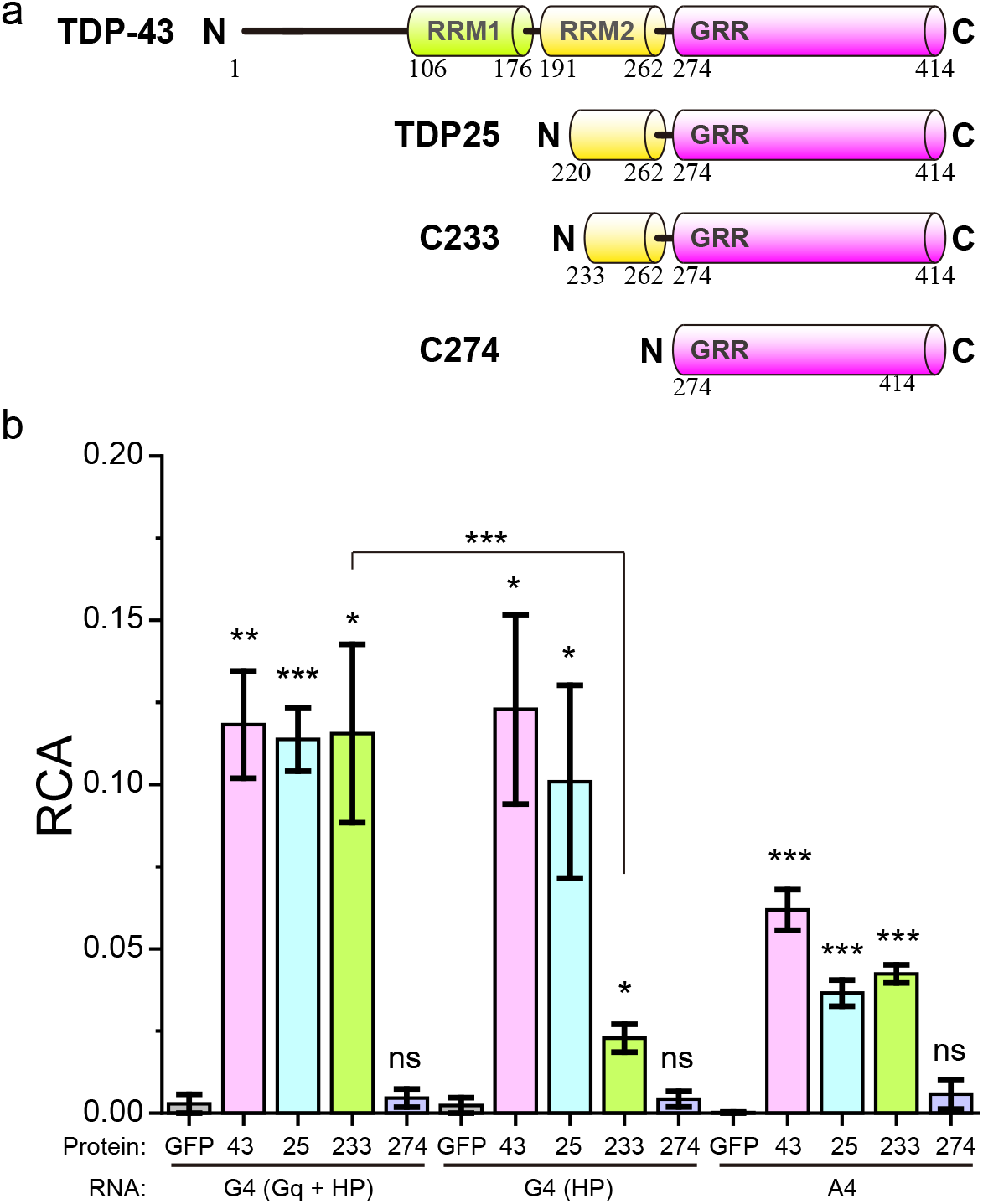
Truncated RRM2 is required for the interaction with G4C2/A4U2-repeat RNA. a) Primary structures of human TDP-43, TDP25, CTF233, and CTF274. Numbers show the amino acid position. b) RCA values when AF-labeled RNAs (G4, and A4) that were folded in 100 mM KCl (Gq + HP) or 100 mM LiCl (HP) were mixed with GFP monomers (GFP), GFP-TDP-43 (43), GFP-TDP25 (25), GFP-CTF233 (233), and GFP-CTF274 (274) with 100 mM KCl (n = 3; mean ± SE). Significance: \**p* < 0.05, \*\**p* < 0.01, \*\*\**p* < 0.001, and ns (*p* ≥ 0.05).

### GGGGCC/AAAAUU hexanucleotide repeats prevent aggregation of TDP25

Next, to demonstrate the activity of G4- and A4-RNA that prevent the aggregation of TDP25, G4 and A4 tagged with bacteriophage-derived M13 sequences (M13-G4 and M13-A4, respectively) were expressed in Neuro2a cells by transfection of plasmid DNA that transcribes such RNAs under the human H1 promoter. The expression of M13-G4, M13-A4, and only M13 tag as a control (M13N) in Neuro2a cells was confirmed by reverse transcription PCR (RT-PCR) (Supplemental Figure S4a). The population of cells expressing M13-G4 and M13-A4 that harbor T25 aggregates in the cytoplasm was decreased, while that of cells expressing M13N was not (Figure 4a). Although the expression levels of T25 in the soluble fraction of the cell lysates were not changed in cells expressing M13-G4, M13-A4, or M13N compared to empty vector transfection, the levels of T25 in the insoluble fraction significantly decreased in cells expressing M13-G4 and M13-A4 (Figure 4b and Supplemental Figure S5a). These results suggest that the expression of G4- and A4-RNA can decrease T25 aggregates in the cell. We reported that an abundant cytoplasmic molecular chaperone HSP70 suppresses T25 aggregation [40]. Although these G4/A4-RNA are supposed to be an aggregation-suppressor for T25, the next question is whether the G4/A4-RNA introduced into cells disturbs the intracellular transcriptome, resulting in the indirect suppression of T25 aggregation through the upregulation of stress-induced proteins, such as heat shock proteins. To this end, transcriptomic analysis was performed using RNA-seq when G4/A4-RNA were expressed in Neuro2a cells. No transcripts with a >2-fold change in the significant expression ratio (*p* < 0.05) were observed in both mRNA and non-coding RNA (ncRNA) (Figure 4c). Furthermore, while adjusting the criteria, four mRNAs (*Znf69, Ppp1ccb, Znf429*, and *Cdon*) of a >1.5-fold change with significance were identified (Supplemental Data); however, these gene-coding proteins were not reported as molecular chaperones. Therefore, it is unlikely that up-regulated chaperone-like genes by G4/A4-RNA expression inhibit T25 aggregation indirectly; in other words, we suggest that these RNAs directly prevent T25 aggregation.

**Figure 4.**
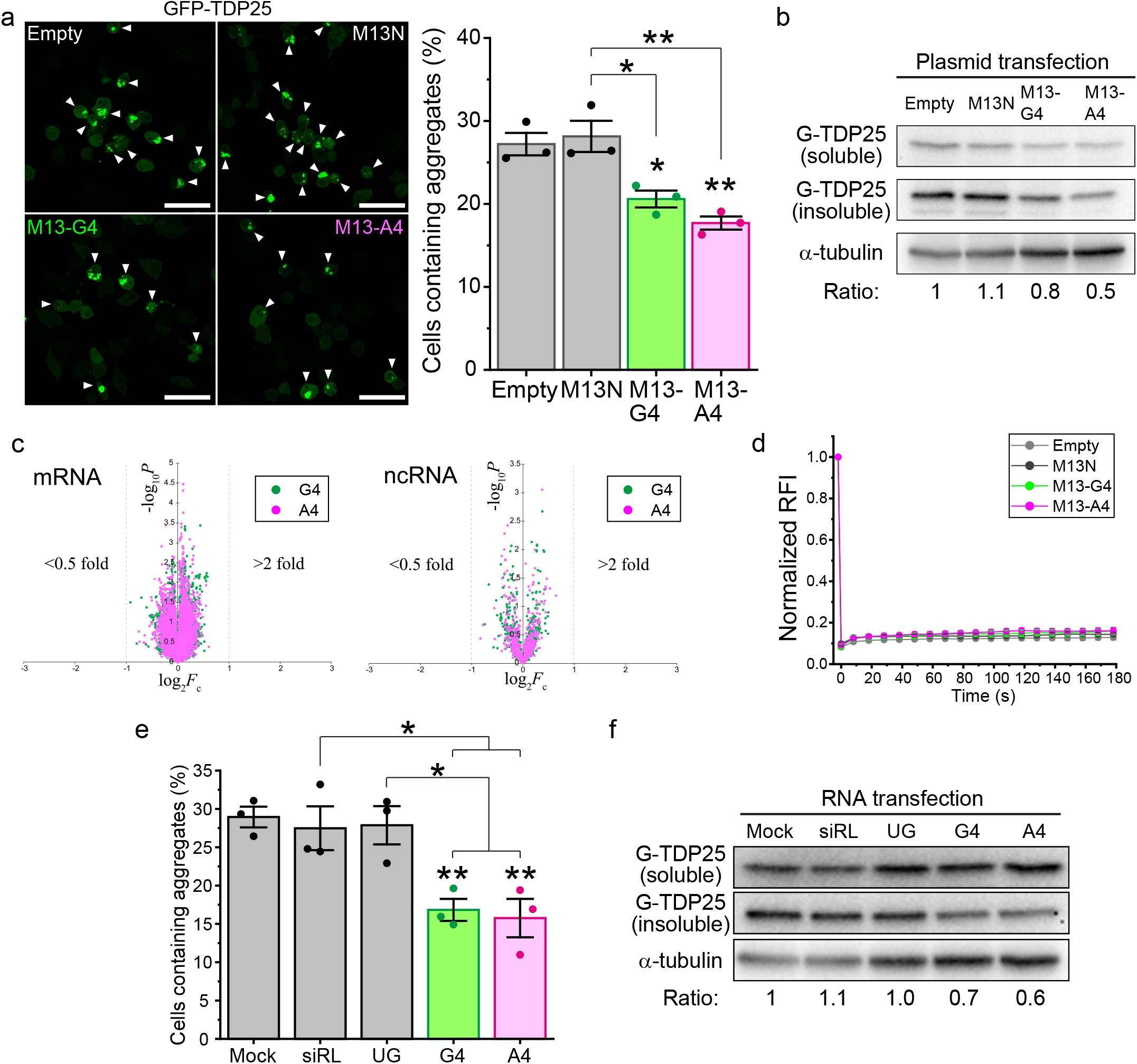
Aggregate formation of GFP-TDP25 in Neuro2a cells expressing G4C2/A4U2-repeat RNA. a) Confocal fluorescence images of GFP-TDP25 (T25) in Neuro2a cells (left). The percentage of cells containing cytoplasmic aggregates of GFP-TDP25 when an empty vector (Empty) or M13-tagged RNA was expressed (M13N, M13-G4, and M13-A4) (right) (n = 3; mean ± SE). Significance: \**p* < 0.05 and \*\**p* < 0.01. b) Western blotting of Neuro2a cells expressing GFP-TDP25 when M13-tagged RNAs were expressed. The value on the bottom indicates the intensity ratio of GFP-TDP25 in the insoluble fraction compared with that in the empty vector transfection. The gel loading the insoluble fraction of the cell lysate stained with Coomassie brilliant blue as an internal control was represented in Supplemental Figure S5a. c) Volcano plot of mRNA and ncRNA expression change when M13-G4 or -A4 was expressed in Neuro2a cells compared with the empty vector transfection. d) Mobile fraction of GFP-TDP25 in the cytoplasmic aggregates determined using FRAP (n = 9; mean ± SE). e) The percentage of cells containing cytoplasmic aggregates of GFP-TDP25 when synthetic RNA (Mock, siRL, UG, G4, or A4) was transfected (n = 3; mean ± SE). Significance: \**p* < 0.05 and \*\**p* < 0.01. f) Western blotting of Neuro2a cells expressing GFP-TDP25 when synthetic RNA (Mock, siRL, UG, G4, or A4) was transfected. The value on the bottom indicates the intensity ratio of GFP-TDP25 in the insoluble fraction compared with that in the Mock RNA transfection. The gel loading the insoluble fraction of the cell lysate stained with Coomassie brilliant blue as an internal control was represented in Supplemental Figure S5b.

Are the biophysical molecular dynamics of T25 within the aggregates then changed in the expression of G4/A4-RNA? To investigate this issue, fluorescence recovery after photobleaching (FRAP) analysis in live Neuro2a cells was performed. T25 was quite immobile and formed amyloids in the aggregates [8,39]; however, the expression of M13-G4/A4 did not alter the immobility of T25 (Figure 3d), suggesting that G4/A4-RNA may act on T25 in the diffusive states in the cytoplasm to prevent aggregation before the formation of deposited aggregates. As a possible reason why G4/A4-RNA cannot act on the deposited aggregates, HSP70 overexpression does not dramatically increase the mobile population of T25 in the aggregates [8,40]. HSP70 is colocalized and dynamically interacts with T25 aggregates [40]; however, RNA is not stained inside the cytoplasmic T25 aggregates [8]. This unremarkable difference in T25 mobility in the aggregates, even when overexpressed HSP70 interacts with the aggregates, is consistent with the no significant changes in T25 mobility in the aggregates when G4/A4-RNA were expressed, as shown here. Therefore, G4/A4-RNA may inhibit the aggregation and assembly process before T25 forms aggregates in the cytoplasm that can be viewed by microscopes. These RNAs would maintain the oligomeric states of T25 and/or prevent its intermolecular assembly.

Next, to demonstrate that G4/A4-RNA without any tag directly prevents T25 aggregation, the aggregate formation of T25 in Neuro2a cells introduced with RNA was analyzed. Transfection of synthetic 24-mer UG repeat RNA (UG) and siRNA against the Renilla luciferase mRNA (siRL) [41] as controls did not change the population of cells containing T25 aggregates in the cytoplasm compared to mock transfection; however, the direct transfection of synthetic G4/A4-RNA significantly decreased the proportion of cells containing T25 aggregates (Figure 3e). Although the expression levels of T25 in the soluble fraction of the cell lysates were not changed in cells expressing M13-G4, M13-A4, or M13N compared to empty vector transfection, the levels of T25 in the insoluble fraction significantly decreased in cells expressing M13-G4 and M13-A4 (Figure 4b). The levels of T25 in the insoluble fraction decreased in G4- and A4-RNA-transfected cells (Figure 4f and Supplemental Figure S5b). These results confirm the hypothesis that G4/A4-RNA can be used as an aggregation inhibitor for T25 by introducing it into cells. Furthermore, this suggests that RNA that binds to TDP-43, as in the case of UG repeat RNA, is not a sufficient condition for the prevention of T25 aggregation.

Because G2-RNA but not A2-RNA interacted with TDP25 (Figure 1d), we next investigate whether G2/A2-RNA can reduce the T25 aggregate. M13-tagged G2/A2-RNA was expressed as much as M13-tagged G4/A4-RNA in Neuro2a cells (Supplemental Figure S6a). G2/A2-RNA expression in Neuro2a cells did not change the population of cells that harbor T25 aggregates and the levels of T25 in the insoluble fraction (Supplemental Figure S6b). G2 is believed to be too short to form the Gq structure efficiently while it is not entirely incapable of forming Gq as intermolecular assemblies. The interaction strength of G2 with T25 was reduced by approximately 60% compared to G4 (Figure 1d). Therefore, for the effective inhibition of the T25 aggregation by r(G4C2) in the cell, a certain degree of interaction strength with T25, which corresponds to that of four repeats of r(G4C2), may be required. Furthermore, the r(A4U2) may also require four repeats to efficiently suppress the T25 aggregation.

### GGGGCC/AAAAUU hexanucleotide repeats prevent aggregation of TDP-43 and ameliorate its cytotoxicity

Although TDP25 is highly prone to aggregation compared to TDP-43 and its CTFs [10,13,39,42], the mislocalization of full-length TDP-43 with its aggregation is thought to be more associated with pathogenicity [14,43,44]. However, wild type TDP-43 is not mislocalized and does not form any cytoplasmic aggregates in Neuro2a cells [8]. Thus, we chose a variant of TDP-43 that lacks the ability of nuclear localization and exporting signal sequences and carries amino acid substitutions of C173S/C175S (TDP43CS) [45]. Furthermore, because GFP-tagged TDP43CS (T43CS) did not form cytoplasmic aggregates in Neuro2a cells (data not shown), we used 293 cells, in which T43CS formed cytoplasmic aggregates efficiently. The expression of M13-G4, M13-A4, and M13N in 293 cells was confirmed using RT-PCR (Supplemental Figure S4b). The population of cells that harbor T43CS aggregates in the cytoplasm was significantly reduced in cells expressing M13-G4 and M13-A4 compared to that of cells expressing M13N and empty vector-transfected cells as controls (Figure 5a). Next, to biochemically detect the reduction in aggregates by expression of M13-G4/A4, these cells were lysed in a buffer containing 0.1% SDS and then fractionated using centrifugation. Almost all T43CS was recovered in the soluble fraction, and no significant change in the T43CS amount both in the soluble and insoluble fraction was observed in G4/A4-RNA-expressed cells (Figure 5b and Supplemental Figure S5c). Therefore, the aggregates of T43CS may be more loosely assembled than those of T25. Furthermore, the filter retardation assay showed that the amount of T43CS trapped on the membrane was decreased by M13-G4/A4 expression (Figure 5b), suggesting that T43CS may not exist as complete monomers and may probably exist as oligomers and/or soluble aggregates that were larger than the pore size of the membrane (>0.2 mm); furthermore, they can be solubilized by SDS, and G4/A4-RNA decreases oligomers and/or soluble aggregates of T43CS likely by affecting its oligomerization process. FRAP analysis revealed that T43CS was also considerably immobile in the aggregates as T25, and the expression of M13-G4/A4 did not alter the immobility of T43CS (Figure 5c), suggesting that G4/A4-RNA may also act on T43CS in the diffusive states in the cytoplasm and likely not in the aggregates deposited. This is consistent with the findings of the western blot and filter retardation assays. Furthermore, because the cytoplasmic mislocalization of TDP-43 is cytotoxic [14], to demonstrate whether G4/A4-RNA expression ameliorates cytotoxicity, we counted rounded and shrinking cells, indicating cytotoxicity and the ongoing progression of cell death in adherent cells [46]. The population of rounded and shrinking cells was dramatically reduced in M13-G4/A4-expressing cells compared to M13N expression and empty vector transfection as controls (Figure 5d), suggesting that G4/A4-RNA plays a role in the reduction of cytotoxicity in T43CS-expressing cells. Therefore, G4/A4-RNA could allow cells with cytoplasmic mislocalization of TDP-43 to be protected with a decrease of its oligomers/soluble aggregates.

**Figure 5.**
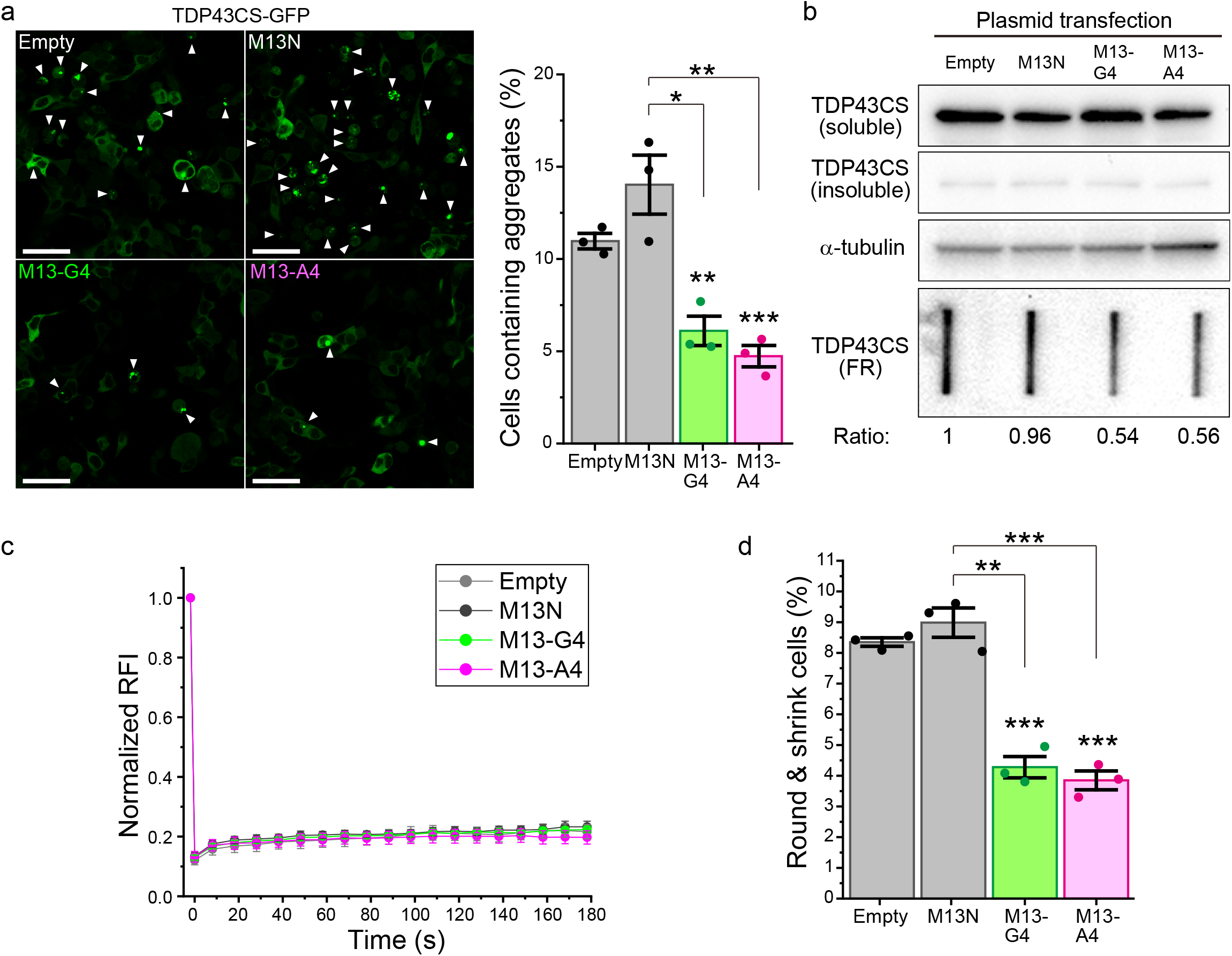
Aggregate formation of GFP-tagged TDP-43CS in 293 cells expressing G4C2/A4U2-repeat RNA. a) Confocal fluorescence images of GFP-tagged TDP-43CS (T43CS) in 293 cells. The white arrows indicate the cells containing the aggregates (left). The percentage of cells containing cytoplasmic aggregates of GFP-tagged TDP-43CS when empty vector (Empty) or M13-tagged RNA was expressed (M13N, M13-G4, and M13-A4) (right) (n = 3; mean ± SE). Significance: \**p* < 0.05, \*\**p* < 0.01, and \*\*\**p* < 0.001. b) Western blotting (*top*) and filter retardation (*bottom*) of 293 cells expressing GFP-tagged TDP-43CS when M13-tagged RNAs were expressed. The value shown on the bottom of the filter retardation (FT) blot images indicates the intensity ratio of the retarded GFP-tagged TDP-43CS on the membrane compared with the empty vector transfection. The gel loading the insoluble fraction of the cell lysate stained with Coomassie brilliant blue as an internal control was represented in Supplemental Figure S5c. c) Mobile fraction of GFP-tagged TDP-43CS in the cytoplasmic aggregates determined using FRAP (n = 7; mean ± SE). d) The population of TDP-43CS-expressing cells with rounded and shrinking shapes (n = 3; mean ± SE). (a & d) Significance: \*\**p* < 0.01 and \*\*\**p* < 0.001.

## Discussion

We demonstrate that G4/A4-RNA has an anti-aggregation activity against not only TDP25 but also TDP-43. For TDP-43 CTFs, truncated RRM2 lacking the binding ability to UG-repeat RNA is involved in the interaction between TDP-43 CTFs and G4/A4-RNA (Figures 1–3 and Supplemental Figures S1–2). Not only electrostatic but also hydrophobic interaction may contribute to the binding between TDP25 and G4/A4-RNA unlike TDP-43 (Figure 1c). These results suggest that G4/A4-RNA likely recognizes not a distinct RNA recognition motif but certain structures during the aggregation process of TDP-43 and its CTFs. Furthermore, because C274 rarely forms the cytoplasmic aggregates [39], oligomers/aggregates would be required for the RNA binding. G4-RNA can fold in a mixed state between Gq and hairpin structure in the presence of potassium ion [30]; however, A4-RNA is predicted to exhibit no optimal secondary structure thermodynamically by an RNAfold prediction server [47], suggesting that specific structures or folding states of RNA would not be necessarily involved in the aggregation-suppressing activity. Otherwise, G4- and A4-RNA would recognize the different structures and oligomeric states of TDP25 and TDP-43 during aggregation, respectively. Furthermore, because UG-repeat RNA did not prevent the aggregation of TDP25, the sequence specificity acting as an aggregation-suppressor is surely required. Because the exogenous introduction of G4/A4-RNA did not affect the intracellular transcriptome (Figure 4c) and ameliorated cellular adhesion properties (Figure 5d), which are depressed by the cytoplasmic expression of TDP-43, these RNAs may recover and probably maintain the proteostasis in the cell by preventing aggregation. Repeat sequences of G4C2/A4U2 have been observed in transcripts using bioinformatic local alignment search tools such as BLAST. The most frequent repeats of them and even of non-pathogenic *C9orf72* transcripts are two. These transcripts that carry repeats of G4C2 or A4U2 would orchestrate the prevention of aggregation of TDP-43 and its CTFs in the cell. As described above, although here we use a short minimum repeat of G4C2 to form the Gq structure intramolecularly, the abundant Gq structure introduced into the cells is possibly a double-edged sword if it is used as a therapeutic drug as it can encourage the abnormal assembly of RNPs such as TDP-43. However, since the A4U2 repeat does not have a specific structure, its four repeats would be safer than the G4C2 repeat. In contrast, because the pathogenic expansion of G4C2 hexanucleotide repeat (>~20) in *C9orf72* transcripts produces toxic aggregates of dipeptide repeats by repeat-associated non-ATG translation [48,49], they would rather be cytotoxic; however, our results show that the non-pathogenic short hexanucleotide repeats may act as an aggregation-suppressor against TDP-43 and TDP25.

In conclusion, this is the first demonstration of aggregation-suppressing RNA against aggregation-prone TDP-43 and TDP25 in the cell. We were able to demonstrate that G4/A4-RNA has anti-aggregation activity against TDP-43 and TDP25. Our study has not clarified whether G4/A4-RNA is a molecular chaperone widely assisting the folding of nascent polypeptides; however, we revealed a new function of RNA, which reduces the cytotoxicity of mislocalized TDP-43 and maintains proteostasis. This report will pave the way for new therapeutic and preventive strategies for ALS/FTD (i.e. such as oligonucleotide therapeutics).

## Materials & Methods

### Construction of plasmids

The plasmids for the expression of GFP-tagged TDP25 (pGFP-TDP25), 233–414 amino acids CTF (C233), and 276–414 amino acids CTF (C274) were used as per the protocol established in previous reports [8,39]. The synthetic DNA fragment for a variant of TDP-43 that lacks nuclear localization and exporting signal sequences and carries amino acid substitutions of C173S/C175S (TDP-43CS-mNLS/NES) [45], was ordered via DNA fragment synthesis services (Eurofins Genomics, Tokyo, Japan). The fragment was inserted into the HindIII and BamHI sites of pmeGFP-N1 (pTDP43CS-GFP).

For purification of GFP-tagged wild type TDP-43, synthetic cDNA fragment for polyhistidine tag and Strep-tag (hereafter, underlined primer sequences denote restriction enzyme sites for cloning) (5’-<u>TGTACA</u>GTCACCATCATCACCATCACTGGAGCCACCCGCAGTTCGAAAAATAA<u>GCGGCCGC</u>-3’) was inserted into the BsrGI and NotI sites of wild type TDP-43 fused with meGFP at the C-terminal, as used in our previous study (pTDP43-GFP-HS) [8]. For purification of GFP-TDP25 and GFP monomers, synthetic cDNA fragment for polyhistidine tag and Strep-tag (5’-<u>GCTAGC</u>CACCATGCACCATCATCACCATCACTGGAGCCACCCGCAGTTCGAAAAACT<u>ACCGGT</u>-3’) was inserted into the NheI and AgeI sites of pGFP-TDP25 and pmeGFP-N1 (pHS-GFP-TDP25 and pHS-GFP, respectively).

The synthetic DNA fragment for M13-tagged RNA (M13N, M13-G4, M13-A4, M13-G2, and M13-A2) (Supplemental Table S1) was ordered using DNA oligonucleotide synthesis services (Thermo Fisher, Waltham, MA, USA). The fragments were inserted into the BglII and HindIII sites of pSUPERIOR.neo (pSUPn-M13N, -M13-G4, -M13-A4, -M13-G2, and -M13-A2).

### RNA synthesis

Synthetic RNAs were ordered using RNA oligonucleotide synthesis services (AJINOMOTO BioPharma, Osaka, Japan). All sequences used in this study are presented in Supplemental Table S2. The 5’ end of the RNAs was labeled with Alexa Fluor 647, a fluorescent dye, followed by purified by high performance liquid chromatography using oligonucleotide synthesis and purification services (AJINOMOTO BioPharma). Fluorescently tagged RNAs were not used for introduction into the cells. The sequence of siRNA against renilla luciferase mRNA (siRL) was obtained from ref.[41].

### Cell culture and transfection

Murine neuroblastoma Neuro-2a cells (#CCL-131; ATCC, Manassas, VA, USA) and Flp-In T-REx 293 cells (Thermo Fisher) were maintained in DMEM (D5796; Sigma-Aldrich, St. Louis, MO, USA) supplemented with 10% FBS (12676029, Thermo Fisher), 100 U/mL penicillin G (Sigma-Aldrich), and 0.1 mg/ml streptomycin (Sigma-Aldrich) as previously described [8]. Neuro2a (2.0 × 10^5^) or Flp-In T-REx 293 cells (4.0 × 10^5^) were spread in a 35 mm glass-bottom dish for microscopic experiments (3910-035; IWAKI-AGC Technoglass, Shizuoka, Japan) or a 35 mm plastic dish for cell lysis (150318; Thermo Fisher) one day before transfection. Before spreading Flp-In T-REx 293 cells, the dishes were coated with 0.01 mg/ml poly-L-lysine. The mixture of pGFP-TDP25 (1 mg) or pTDP43CS-GFP (1 mg) and pSUPERIOPR.neo (1 mg), pSUPn-M13N (0.25 mg mixed with 0.75 mg of the empty vector), pSUPn-M13G4 (1 mg), or pSUPn-M13A4 (1 mg) were transfected using 5.0 ml of Lipofectamine 2000 (Thermo Fisher). After overnight incubation for transfection, the medium was replaced. Neuro2a and Flp-In T-REx 293 cells were cultured for 24 and 48 h after transfection, respectively, before microscopic analysis or cell lysis. For protein purification, Neuro2a cells (3.0 × 10^6^) were spread in a 150 mm glass-bottom dish (TR4003; NIPPON Genetics, Tokyo, Japan) 1 d before transfection. A mixture of 14 mg of salmon sperm DNA (BioDynamics Laboratory Inc., Tokyo, Japan) and 16 mg of expression plasmid DNA (pTDP-43-GFP-HS, pHS-GFP-TDP25, or pHS-GFP) was diluted in a 150 mM NaCl solution, followed by the addition of 77.5 mg polyethyleneimine (PEI) in a 10 mM Tris-HCl buffer (pH 8.0). The mixture was added to the dishes after 15 min of incubation. After 24 h of incubation, the cells were lysed for protein purification. For purification, 30, 30, and 2 dishes were prepared for TDP-43, TDP25, and GFP monomers, respectively. For sequential transfection of RNA and plasmids, Neuro2a cells (2.0 × 10^5^) were spread in a 35 mm glass-bottom dish (IWAKI-AGC Technoglass) one day before RNA transfection. After the cells were transfected with RNA (25 pmol) using 5.0 mL of Lipofectamine RNAi MAX (Thermo Fisher), the cells were incubated for 7 h. After the medium exchange, the cells were transfected with pGFP-TDP25 (1 mg) using 2.5 ml of Lipofectamine 2000 (Thermo Fisher Scientific). The cells were cultured for 24 h after plasmid transfection before the microscopic analysis.

### Protein purification

GFP-tagged protein-expressing Neuro2a cells were collected in tubes after trypsinization. Lysis buffer containing 50 mM HEPES-KOH (pH 7.5), 150 mM NaCl, 1% Noidet P-40, and 1× protease inhibitor cocktail (P8340, Sigma-Aldrich) was added to a tube placed on ice, and the supernatant was recovered after centrifugation at 20,400 × *g* for 10 min at 4°C. Ni-NTA agarose beads (25 mL slurry for GFP monomers and 75 mL slurry for TDP-43 and TDP25; Fujifilm Wako Chemicals, Osaka, Japan) were added to the supernatant, followed by rotation and incubation for 60 min. The beads were washed thrice in buffer containing 50 mM HEPES-KOH (pH 7.5), 150 mM NaCl, 20 mM imidazole, and 0.2% Noidet P-40. After the proteins were eluted in a buffer containing 50 mM HEPES-KOH (pH 7.5), 150 mM NaCl, 250 mM imidazole, and 0.2% Noidet P-40, the protein solutions were applied to a spin column containing Strep-Tactin XT Superflow suspension (Zymo Research, Irvine, CA, USA). After washing the column in a buffer containing 50 mM HEPES-KOH (pH 7.5), 150 mM NaCl, and 0.2% Noidet P-40, the proteins were eluted in a buffer containing 50 mM HEPES-KOH (pH 7.5), 150 mM NaCl, 0.2% Noidet P-40, and 50 mM biotin (Fujifilm Wako Chemicals). The proteins were concentrated in a buffer containing 20 mM HEPES-KOH (pH 7.5) and 0.2% Noidet P-40 using a dialysis spin column (Amicon Ultra-10; Merck, Darmstadt, Germany). The concentrations of purified fluorescent proteins were measured using fluorescence correlation spectroscopy (FCS). The purified proteins were immediately used for FCCS measurements. For mass spectroscopic analysis, the purified proteins were separated on a 5-20% gradient gel (ATTO, Tokyo, Japan) using SDS-PAGE, followed by silver staining using silver stain reagents (423413; Cosmo Bio, Tokyo, Japan). The bands of interest were cut and analyzed using MALDI-TOF-MS, followed by peptide mass fingerprinting that was performed using an outsourced service (Genomine, Pohang, South Korea).

### Fluorescence cross-correlation spectroscopy

Before FCCS measurement, 10 mM Alexa Fluor 647-labeled RNAs were heat-denatured and cooled (7°C/min) in a buffer containing 20 mM Tris-HCl (pH 8.0) and 100 mM KCl for the mixture of G-quadruplex and hairpin or LiCl for the only hairpin. Neuro2a cells were lysed in a buffer containing 20 mM HEPES-KOH (pH 7.5), 1% Noidet P-40, and 1× protease inhibitor cocktail (Sigma-Aldrich). The supernatants of the lysates were recovered after centrifugation at 20,400 × *g* for 10 min at 4°C. The supernatant was mixed with 100 nM RNA and 100, 150, 200, or 250 mM KCl and incubated for five minutes before the measurement. FCCS was performed using LSM 510 META + ConfoCor3 (Carl Zeiss) with a C-Apochromat 40×/1.2NA W. UV-VIS-IR water immersion objective lens (Carl Zeiss) as previously reported [8]. The photons were recorded for 100 s. Curve fitting analysis for the acquired autocorrelation function was performed using ZEN software (Carl Zeiss) with a model for two-component 3D diffusion involving a one-component exponential blinking fraction, as shown in the following equation (Eq.) 1.

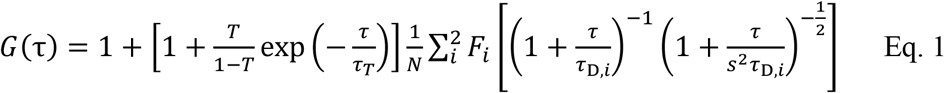

where G(t) represents an auto- or cross-correlation function of the time interval t; t*_D, i_* represents the diffusion time (*i* = 1 or 2); *F_i_* represents the fraction of the component (*i* = 1 or 2); *N* represents the average number of particles in the confocal detection volume; *T* and t*_T_* represent the exponential blinking fraction and its meantime, respectively (these are zero for cross-correlation function); *s* represent the structural parameter determined by analyzing the standard fluorescent dyes (ATTO 488 and Cy5) on the same day. The relative cross-correlation amplitude (RCA) was calculated using Eq. 2.

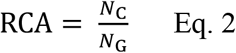

where *N*_C_ and *N*_G_ represent the average number of interacting and green molecules, respectively.

### RT-PCR

Total RNA of cells expressing M13-tagged RNA was isolated using an RNeasy Mini Kit (QIAGEN, Venlo, Netherlands) according to the manufacturer’s instructions. The RNA was eluted in diethyl pyrocarbonate-treated ultrapure water. To detect M13-tagged RNAs, total RNA (1.9 mg) was used to synthesize first-strand cDNA using the Mir-X miRNA First-Strand Synthesis Kit (TaKaRa) according to the manufacturer’s instructions. PCR was performed using a 0.25 U PrimeSTAR HS DNA polymerase (TaKaRa) with 0.2 mM dNTP mixture, 2 pmol of M13-forward primers (5’-gtaaaacgacggccagt-3’), 2 pmol of M13-reverse primers (5’-gagcggataacaatttcacacagg-3’), and template cDNA (corresponding to 15 ng of total RNA). Amplification was performed using a PC320 thermal cycler (Astec-Bio, Fukuoka, Japan). The products were separated on a 5–20% polyacrylamide gel (ATTO) in Tris-glycine buffer. The gel was stained with SYBR Gold (Thermo Fisher Scientific), and gel images were acquired using a Limited-STAGE II gel imager with blue LED light (AMZ System Science, Osaka, Japan).

### Confocal microscopy and cell counting

GFP-expressing cells were observed using an LSM 510 META microscope (Carl Zeiss, Jena, Germany) with a Plan-Apochromat 10×/0.45NA objective (Carl Zeiss). GFP was excited at a wavelength of 488 nm. The excitation beam was divided using an HFT405/488. Fluorescence was corrected via a 505-570 nm band-pass filter (BP505-570) and then acquired using a photo-multiplier tube. The pinhole was set to 72 mm. Transmission bright-field images were acquired simultaneously with GFP fluorescence images. Aggregate-positive cells were counted using the Fiji-ImageJ software. The population of cells that harbored cytoplasmic aggregates was normalized to the number of GFP-positive cells. Multiple fluorescent images were acquired at randomly selected locations and then GFP-positive and aggregate-positive cells were counted using the all acquired images. Aggregate-positive cells were counted when the brightness of the structures was higher than that in the cytoplasm. The total number of GFP-positive cells was counted ranged from 200 to 400 per trial. Statistical analysis of three independent trials was performed, and the results are plotted in the graph (n = 3; mean ± SE). The population of rounded and shrinking cells was normalized to the total number of cells in the bright field image. The total number of cells was counted ranged from 280 to 400 per trial from the randomly acquired multiple bright field images. Statistical analysis of three independent trials was performed, and the results are plotted in the graph (n = 3; mean ± SE).

### Fluorescence recovery after photobleaching

Photobleaching experiments were performed with an LSM 510 META system using a C-Apochromat 40×/1.2NA W Korr. UV-VIS-IR M27 objective lens (Carl Zeiss), as previously reported [8,39,40]. GFP was excited and photobleached at 488 nm. The GFP fluorescence was collected through a bandpass filter (BP505-570). The pinhole size was set to 200 μm, the zoom factor was set to 5×, and the interval time for image acquisition was set to 10 s. The X- and Y-scanning sizes were 512 pixels, and the GFP was photobleached at 488 nm with 100% transmission for less than 1.92 s. Relative fluorescence intensities (RFIs) were measured using the Fiji-ImageJ software and calculated as reported previously [8,39,40]. Normalized relative fluorescence intensities (nRFIs) were calculated using the mean RFI of GFP-TDP25 in the aggregates with an empty vector transfection just after photobleaching.

### Cell lysis, solubility assay, and filter retardation assay

Cells expressing GFP-tagged TDP25 and TDP-43CS were lysed and fractionated as reported previously [8]. Briefly, cells were lysed in a lysis buffer containing 25 mM HEPES-KOH (pH 7.5), 150 mM NaCl, 1% Noidet P-40, 0.1% SDS, 0.25 U/mL of benzonase (Sigma-Aldrich), and 1× protease inhibitor cocktail (Sigma-Aldrich). After centrifugation (20,400 × *g*, 10 min, 4°C), the supernatants were recovered, and the pellets were solubilized in PBS containing 1 M urea. SDS-PAGE sample buffer-mixed lysates were denatured at 98°C for 5 min. The samples were applied to a 12.5% polyacrylamide gel (ATTO) and subjected to electrophoresis in an SDS-containing buffer. Proteins were transferred onto polyvinylidene difluoride membranes (Cytiva, Marlborough, MA, USA). The membrane was blocked with PBS-T buffer containing 5% skim milk for 1 h. Horseradish peroxidase-conjugated anti-GFP (#598-7, MBL, Tokyo, Japan) or anti-a-tubulin antibodies (#HRP-66031, Proteintech, Rosemont, IL, USA) were reacted in CanGet Signal Immunoreaction Enhancer Solution 1 (TOYOBO, Osaka, Japan). Chemiluminescent signals were acquired using a ChemiDoc MP imager (Bio-Rad, Hercules, CA, USA). As an internal control for the insoluble fraction, the insoluble fractions were applied to a 12.5% polyacrylamide gel (ATTO) and subjected to electrophoresis in an SDS-containing buffer, followed by staining with Coomassie brilliant blue R-250 (Fujifilm Wako Chemicals). Images of the stained gels were obtained using a Limited-STAGE II gel imager with transparency white light (AMZ System Science). For filter retardation assay, the cells were lysed in a PBS buffer containing 1% SDS and 0.25 U/mL of benzonase (Sigma-Aldrich). The lysates were applied to a cellulose acetate membrane with a pore size of 0.2 mm (Advantec Toyo, Tokyo, Japan) using a Bio-Dot SF blotter (Bio-Rad). The membrane was blocked in PBS-T buffer containing 5% skim milk for 1 h. The GFP on the membrane was detected by western blotting, as described above. All the intensities were measured using Fiji-ImageJ. After subtracting the values in the background region where there are no bands, the ratio value of the intensity of TDP-43/TDP25 in the insoluble fraction or trapped on the filter to the control was calculated.

### RNA-seq and data processing

Total RNA of the cells expressing M13-tagged RNA was recovered in the same manner as that of RT-PCR. The total RNA concentration was measured using a Quantus Fluorometer with a QuantiFluor RNA system (Promega). The quality of total RNA was examined using a fragment analyzer system with an Agilent HS RNA kit (Agilent Technologies). Libraries were prepared using an MGIEasy RNA Directional Library Prep Set (MGI Tech Co., Ltd.) according to the manufacturer’s instructions. The concentration of the libraries was measured using a Qubit 3.0 Fluorometer with a dsDNA HS assay kit (Thermo Fisher). The quality of the libraries was analyzed using a fragment analyzer with a dsDNA 915 reagent kit (Advanced Analytical Technologies). Circular DNA was created using the libraries and an MGIEasy Circularization kit (MGI Tech Co., Ltd.), according to the manufacturer’s protocol. DNA Nano Ball (DNB) was prepared using a DNBSEQ-G400RS high-throughput sequencing kit (MGI Tech Co., Ltd.) according to the manufacturer’s protocol. DNBs were sequenced using a DEBSEQ-G400 sequencer (MGI Tech Co., Ltd.). After removing the adaptor sequence using Cutdapt ver. 1.9.1 software, Pair reads that were less than 40 bases and a quality score of less than 20 were removed using a sickle ver. 1.33 software. Mapping was performed by referring to a murine genome reference (GRCm39) using the hisat2 ver. 2.2.1 software. The number of reads was counted using the featureCounts ver. 2.0.0 software. Reads per kilobase per million (RPKM) and transcripts per million (TPM) values were calculated. After removing transcripts with low read counts (< 19), TPM values derived from total RNA obtained four times, independently, were averaged. The mean fold change and *p*-values were calculated for cells expressing M13-G4/A4 in comparison with those of empty vector-transfected cells. Volcano plots were prepared using Microsoft Excel.

## Acknowledgments

A.F. was supported by the Support for Pioneering Research Initiated by the Next Generation (SPRING) program by the Japan Science and Technology Agency (JST) in Hokkaido University (JPMJSP2119). M. K. was partially supported by a Japan Society for Promotion of Science (JSPS) Grant-in-Aid for Scientific Research (B) (22H02578) and Grant-in-Aid for Challenging Research (Exploratory) (22K19886). A.K. was supported by grants from Japan Agency for Medical Research and Development (AMED) (JP22gm6410028 and JP22ym0126808); by grants from JSPS Grant-in-Aid for Scientific Research (22H04826; 18K06201; and 16KK0156); by a grant from Hokkaido University Office for Developing Future Research Leaders (L-Station); by a grant from Hoansha Foundation; by a grant from Hagiwara Foundation of Japan; and by a grant from Nakatani Foundation. We would like to thank Editage (www.editage.com) for English language editing.

## Author contributions

A.F. performed the experiments, analyzed the data, and edited the manuscript. M.K. contributed to the FCCS measurement and its analysis. A.K. conceptualized and supervised this project, performed the experiments, analyzed the data, visualized the data, provided resources, wrote the original manuscript draft, and edited the manuscript.

