## Supplemental Figures & Tables for "Short repeat RNA reduces cytotoxicity by preventing the aggregation of TDP-43 and its 25 kDa carboxy-terminal fragment"

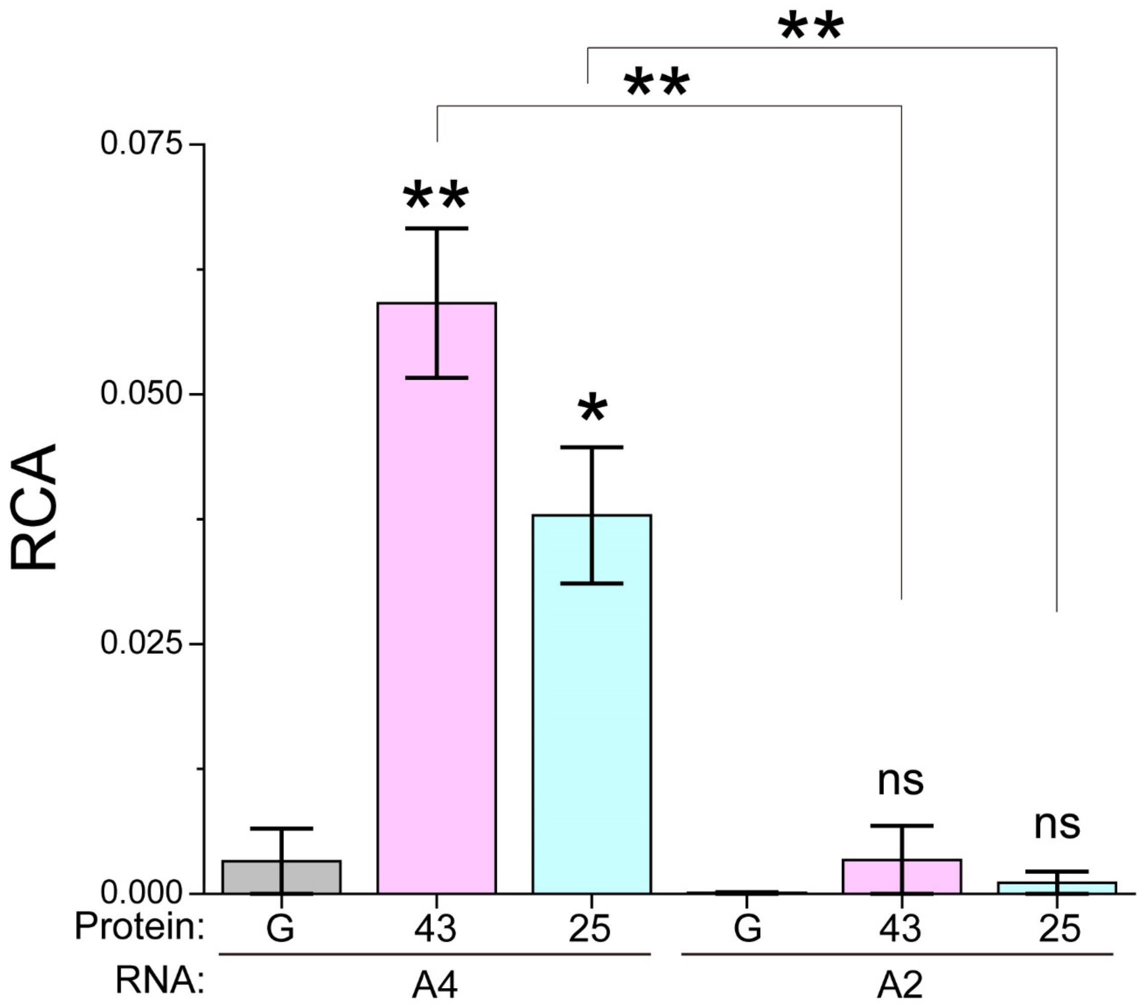


**Supplemental Figure S1. Comparison of RCA values between A4 and A2.**
RCA values when AF-labeled RNAs (A4, and A2) were mixed with GFP monomers (G), GFP-TDP-43 (43), and GFP-TDP25 with 100 mM KCl (n = 3; mean ± SE). Significance: *$p < 0.05$, **$p < 0.01$, and ns ($p \geq 0.05$). The RCA values for A4 and A2 were the same as in Figures 1b & 1d.

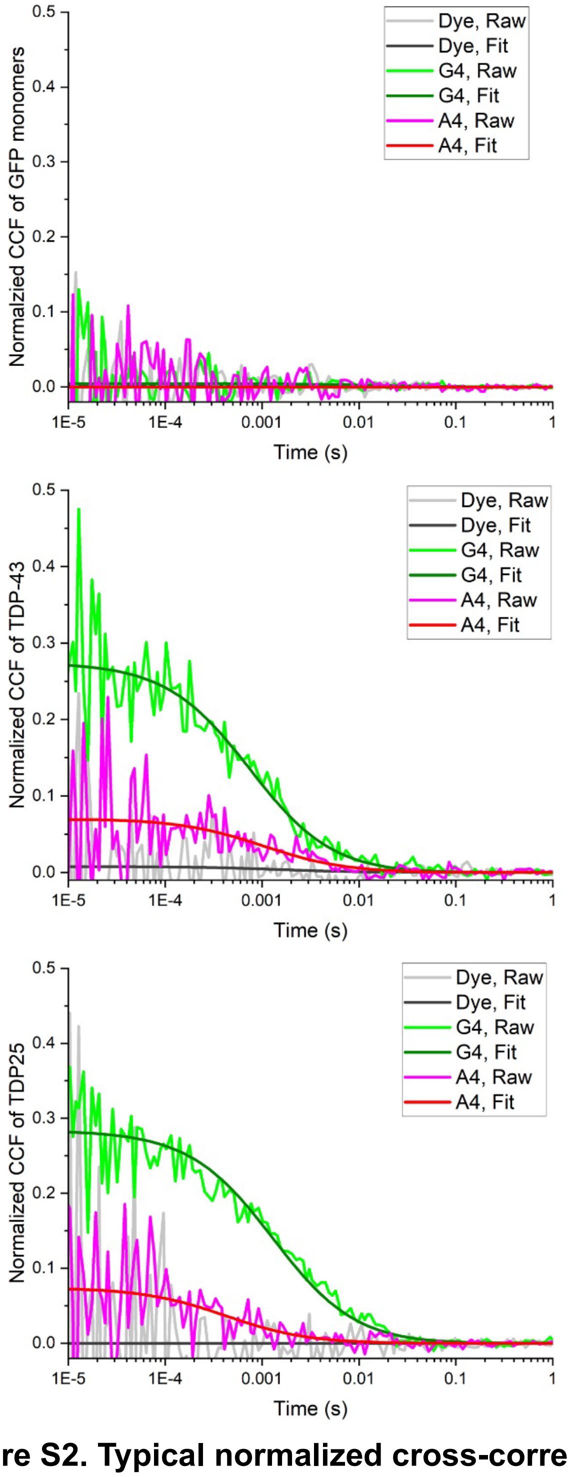


**Supplemental Figure S2. Typical normalized cross-correlation functions (CCFs) when purified proteins were mixed with Alexa Fluor647-labeled RNA.** The purified GFP monomers (*top*), GFP-TDP-43 (*middle*), and GFP-TDP25 (*bottom*) were mixed with Alexa Fluor647 (Dye) or Alexa Fluor647-labeled G4 or A4. Raw CCFs (Raw) and fitted curves (Fit) were represented.No positive RCA between HSP70-GFP and G4 and A4 RNA.

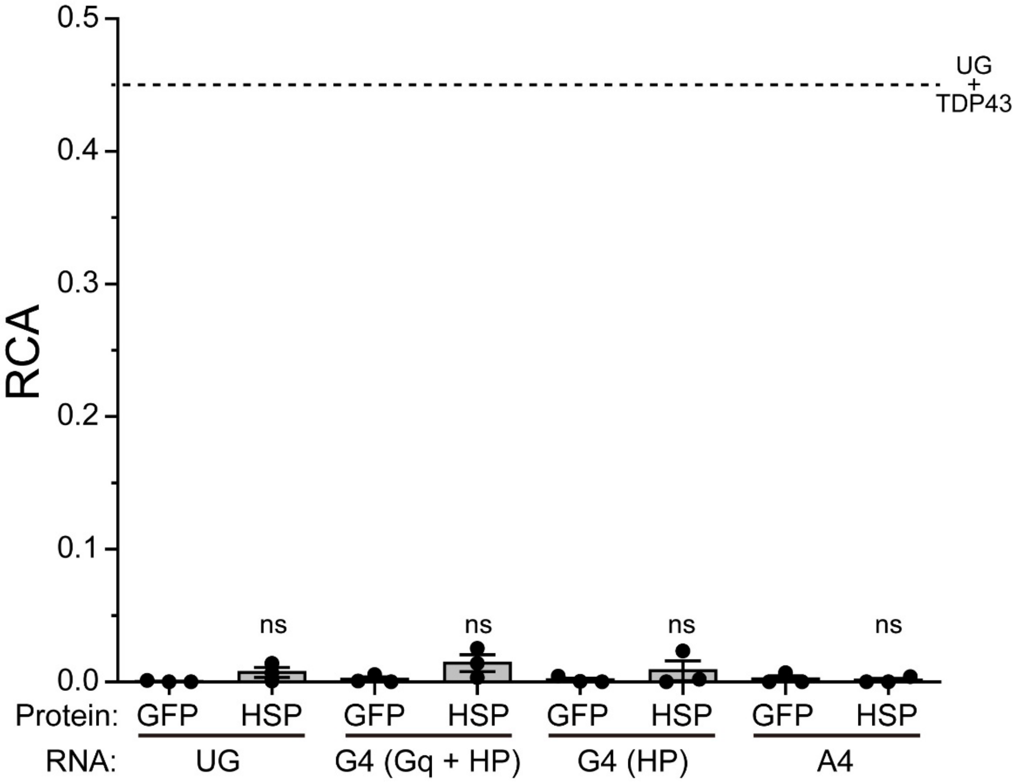


**Supplemental Figure S3. No positive RCA between HSP70-GFP and G4 and A4 RNA.**

RCA values when Alexa Fulor 647-labeled RNAs (UG, G4, and A4) that were folded in 100 mM KCl (Gq + HP) or 100 mM LiCl (HP) were mixed in lysates of Neuro2a cells expressing GFP monomers (GFP) or GFP-tagged HSP70 (HSP) with 100 mM KCl (n = 3; dots and bars indicate independent trials and mean ± SE). Significance: ns ($p \geq 0.05$). The dot line indicates the mean RCA values when UG was mixed with GFP-tagged TDP43 as same as the value in Figures 1b.

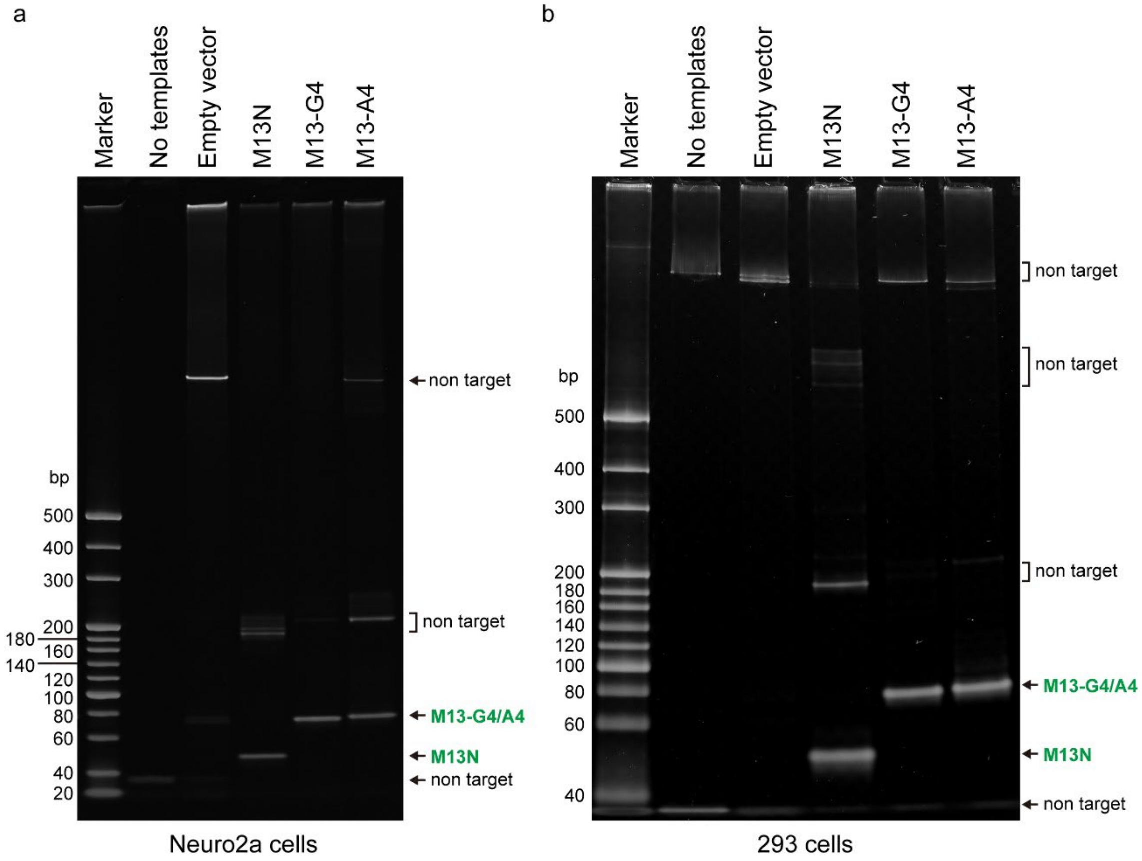


**Supplemental Figure S4. Confirmation of the expression levels of M13-tagged RNA using RT-PCR.** The amplified DNA was analyzed on the gel derived from Neuro2a (left) and Flp-In T-Rex 293 cells (right). The green letters indicate the bands of interest (M13N, M13-G4 and -A4). The values shown on the left of the gel images indicate the size of the molecular weight markers for double-stranded DNA.

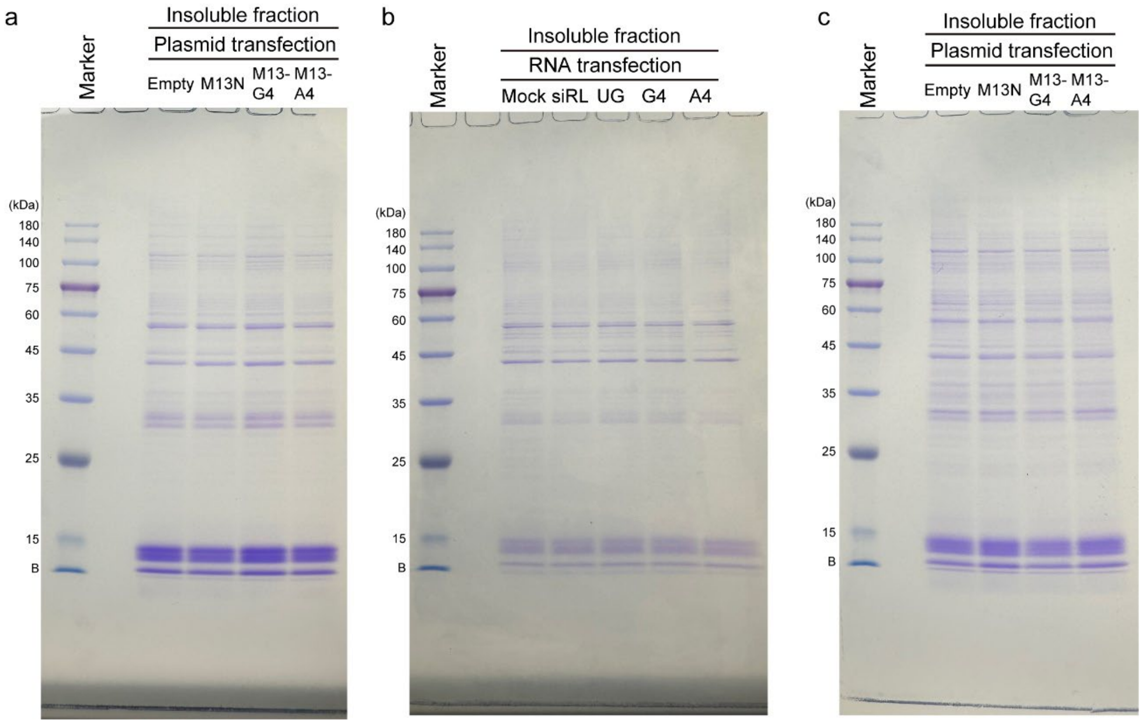


**Figure S5. Loading control of insoluble fraction of cell lysate.**

(a-c) SDS-PAGE gel images of insoluble fraction of cell lysates represented in Fig. 4b (a), 4f (b), and 5b (c) stained with Coomassie brilliant blue R-250. The numbers at the left side of the gel images show the molecular weight of the protein marker standard (Loaded in the lane for 'Marker'), and 'B' indicates the bottom position of the electrophoresis. a) and c) the insoluble fraction of Neuro2a cells expressing GFP-tagged TDP25 (a) and Flp-In T-Rex 293 cells expressing GFP-tagged TDP-43 (c) when an empty vector (Empty) or M13-tagged RNA was expressed (M13N, M13-G4, and M13-A4). b) the insoluble fraction of Neuro2a cells expressing GFP-tagged TDP25 when synthetic RNA (Mock, siRL, UG, G4, or A4) was transfected.

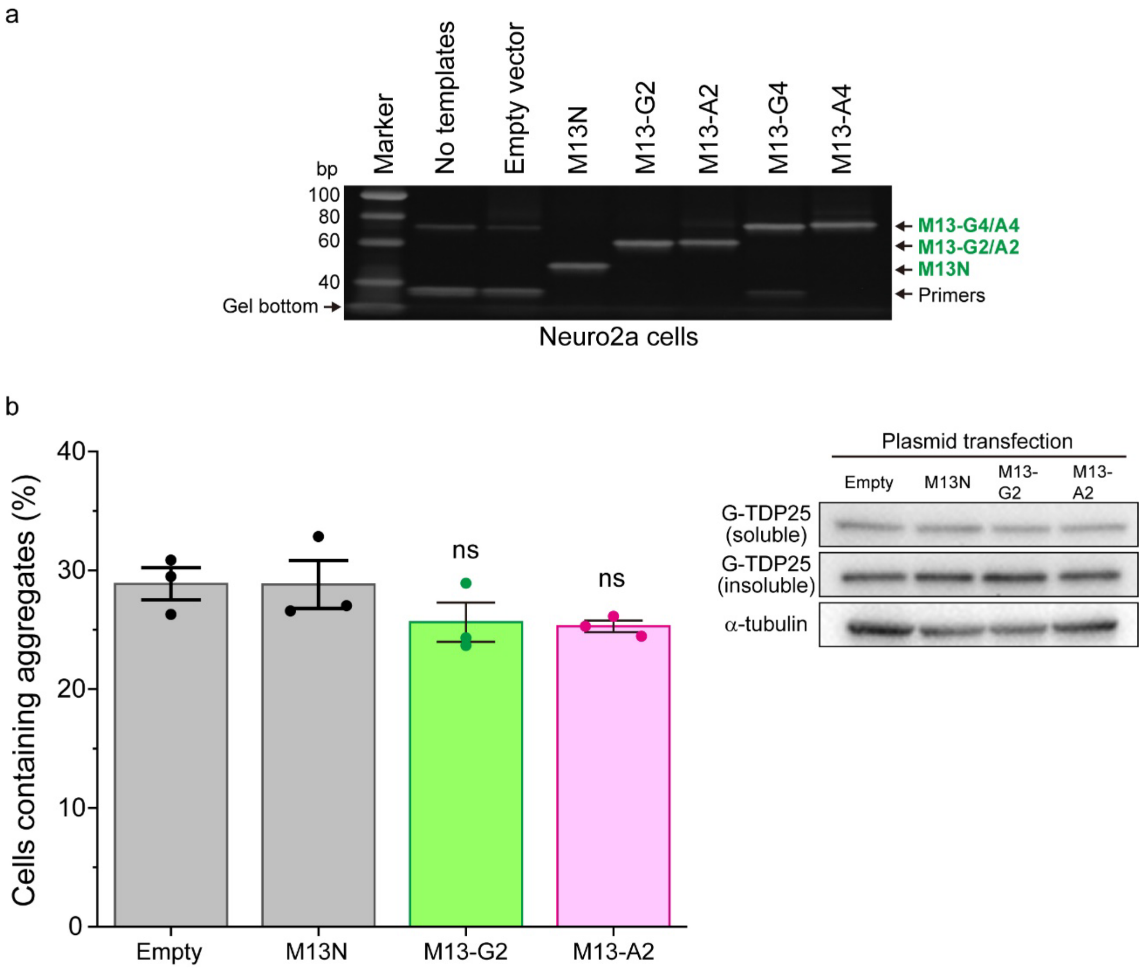


**Supplemental Figure S6. Aggregate formation of GFP-tagged TDP-25 in Neuro2a cells expressing two repeats of G4C2/A4U2-RNA.** a) Confirmation of the expression levels of M13-tagged two or four repeats of r(G4C2) and r(A4U2) (M13-G2, -A2, -G4, and -A4, respectively). M13N is a tag control. Green letters indicate the bands of interest (M13N, M13-G4, and -A4). Unreacted/remained primers were indicated using arrows. b) The percentage of cells containing cytoplasmic aggregates of GFP-TDP25 when an empty vector (empty) or M13-tagged RNA was expressed (M13N, M13-G2, and M13-A2) (*left*) (n = 3; mean ± SE). ns indicates not significant ($p \geq 0.05$). Western blotting of Neuro2a cells expressing GFP-TDP25 when M13-tagged RNAs were expressed (*right*).

### Supplemental Table S1. cDNA sequence for plasmid DNA

| Name | cDNA sequence (5' to 3') |
|---|---|
| **TDP-43CS-mNLS/mNES** | AAGCTTATGTCTGAATATATTCGGGTAACCGAAGATGAGAACGATGAGCCCATTGAAATACCATCGGAAGACGATGGGACGGTGCTGCTCTC<br>CACGGTTACAGCCCAGTTTCCAGGGGCGTGTGGGCTTCGCTACAGGAATCCAGTGTCTCAGTGTATGAGAGGTGTCCGGCTGGTAGAAGGAA<br>TTCTGCATGCCCCAGATGCTGGCTGGGGAAATCTGGTGTATGTTGTCAACTATCCAAAAGATAACGCCGCTGCCATGGATGAGACAGATGCT<br>TCATCAGCAGTGGCCGTGGCCGCTGCAGTCCAGAAAACATCCGATTTAATAGTGTTGGGTCTCCCATGGAAAACAACCGAACAGGACCTGAA<br>AGAGTATTTTAGTACCTTTGGAGAAGTTCTTATGGTGCAGGTCAAGAAAGATCTTAAGACTGGTCATTCAAAGGGGTTTGGCTTTGTTCGTT<br>TTACGGAATATGAAACACAAGTGAAAGTAATGTCACAGCGACATATGATAGATGGACGATGGTCCGACTCCAAACTTCCTAATTCTAAGCAA<br>AGCCAAGATGAGCCTTTGAGAAGCAGAAAAGTGTTTGTGGGGCGCTGTACAGAGGACATGACTGAGGATGAGCTGCGGGAGTTCTTCTCTCA<br>GTACGGGGATGTGATGGATGTCTTCATCCCCAAGCCATTCAGGGCCTTTGCCTTTGTTACATTTGCAGATGATCAGGCCGCGCAGTCTGCCT<br>GTGGAGAGGACGCCGCTGCCAAAGGAATCAGCGTTCATATATCCAATGCCGAACCTAAGCACAATAGCAATAGACAGTTAGAAAGAAGTGGA<br>AGATTTGGTGGTAATCCAGGTGGCTTTGGGAATCAGGGTGGATTTGGTAATAGCAGAGGGGGTGGAGCTGGTTTGGGAAACAATCAAGGTAG<br>TAATATGGGTGGTGGGATGAACTTTGGTGCGTTCAGCATTAATCCAGCCATGATGGCTGCCGCCCAGGCAGCACTACAGAGCAGTTGGGGTA<br>TGATGGGCATGTTAGCCAGCCAGCAGAACCAGTCAGGCCCATCGGGTAATAACCAAAACCAAGGCAACATGCAGAGGGAGCCAAACCAGGCC<br>TTCGGTTCTGGAAATAACTCTTATAGTGGCTCTAATTCTGGTGCAGCAATTGGTTGGGGATCAGCATCCAATGCAGGGTCGGGCAGTGGTTT<br>TAATGGAGGCTTTGGCTCAAGCATGGATTCTAAGTCTTCTGGCTGGGGAATGAATCACGCGGATCC |
| **M13N** | GGATCCCCGTAAAACGACGGCCAGTGAGCTCACCTGTGTGAAATTGTTATCCGCTCTTTTTAAGCTT |
| **M13-G4** | GGATCCCCGTAAAACGACGGCCAGTGGGGCCGGGGCCGGGGCCGGGGCCGAGCTCACCTGTGTGAAATTGTTATCCGCTCTTTTTAAGCTT |
| **M13-A4** | GGATCCCCGTAAAACGACGGCCAGTAAAATTAAAATTAAAATTAAAATTGAGCTCACCTGTGTGAAATTGTTATCCGCTCTTTTTAAGCTT |

**Supplemental Table S2. Synthetic RNA sequence**

| Name | Synthetic RNA sequence (5' to 3') |
|---|---|
| UG | UGUGUGUGUGUGUGUGUGUGUGUG |
| G4 | GGGGCCGGGGCCGGGGCCGGGGCC |
| A4 | AAAAUUAAAAUUAAAAUUAAAAUU |
| G2 | GGGGCCGGGGCC |
| A2 | AAAAUUAAAAUU |
| siRL sense | GCCAAGAAGUUUCCUAAUATT |
| siRL antisense | UAUUAGGAAACUUCUUGGCTT |
